# De novo identification of CD4+ T cell epitopes

**DOI:** 10.1101/2022.11.21.517373

**Authors:** Paul Zdinak, Stephanie Grebinoski, Jessica Torrey, Eduardo Zarate-Martinez, Louise Hicks, Rashi Ranjan, Nishtha Trivedi, Sanya Arshad, Mark Anderson, Dario AA Vignali, Alok V. Joglekar

## Abstract

CD4+ T cells recognize peptide antigens presented on class II Major Histocompatibility Complex (MHC-II) molecules to carry out their function. The remarkable diversity of T cell receptor (TCR) sequences and lack of antigen discovery approaches for MHC-II make profiling the specificities of CD4+ T cells challenging. We have expanded our platform of Signaling and Antigen-presenting Bifunctional Receptors to encode MHC-II molecules presenting covalently linked peptides (SABR-IIs) for CD4+ cell antigen discovery. SABR-IIs can present epitopes to CD4+ T cells and induce signaling upon their recognition, allowing a readable output. Here, we demonstrate that SABR-IIs libraries presenting endogenous and post-translationally modified epitopes can be used for antigen discovery. Using SABR-II libraries in conjunction with single cell RNA sequencing, we de-convoluted multiple highly expanded TCRs from pancreatic islets of Non-Obese Diabetic (NOD) mice. We compounded antigen discovery by incorporating computational TCR similarity prediction metrics followed by experimental validation. Finally, we showed SABR-IIs presenting epitopes in class II HLA alleles can be used for antigen discovery for human CD4+ T cells. Taken together, we have developed a rapid, flexible, scalable, and versatile approach for the de novo identification of CD4+ T cell ligands from single cell RNA sequencing data using experimental and computational approaches.

## INTRODUCTION

A hallmark of the adaptive immune system is the ability to raise antigen-specific responses. This is accomplished for αβT cells through the T cell receptor (TCR), which comprises TCRα and TCRβ chains^1^. TCRs from CD4+ T cells recognize peptide epitopes on MHC-II or human leukocyte antigen (HLA-II) in humans. The estimated size of the mature TCR repertoire is 10^8^-10^10^ unique TCRs in mice and 10^9^-10^12^ unique TCRs in humans^2-4^. Recognition of foreign antigens such as those from SARS-CoV-2 and tumor neoantigens by CD4+ T cells leads to their protective function^5,6^. On the other hand, recognition of self-antigens such as Insulin in Type 1 Diabetes, leads to pathogenic CD4+ T cell responses^7,8^. Furthermore, regulatory T cells (Treg) can bind to self-antigens and prevent autoimmunity in peripheral sites, including within tissues^9^. The specificity of CD4+ T cells is key to their function, highlighting a need for antigen discovery approaches tailored for MHC-II and HLA-II^10^.

Traditionally, antigen-specific CD4+ T cells have been studied using functional assays that measure proliferation, cytokine release, or cytotoxicity in response to stimulation with synthetic peptides^11-14^. These assays are sensitive but inherently lack throughput and are limited to investigating 10s of peptides simultaneously. Techniques such as barcoded tetramers can efficiently detect T antigen-specific T cells but are limited to the interrogation of 100s of specificities simultaneously^15-20^. Approaches using peptide-MHC-II (pMHC-II) multimers are further limited by the instability of multimers and lower affinities of CD4^+^ TCRs, precluding the ability to survey important populations of antigen-specific CD4^+^ T cells^21,22^. Unbiased approaches such as yeast display and combinatorial peptide libraries have been used to identify epitopes *de novo*, but these methods often identify non-physiological epitopes (altered peptide ligands or mimotopes) without the ability to preferentially encode specific epitopes, are highly laborious, and in the case of yeast display, rely on soluble TCR generation^23-26^. Recently described cell-based methods are emerging approaches for TCR-directed antigen discovery. These methods preserve physiological TCR-pMHC interactions, are capable of presenting large and defined epitope libraries, and do not require significant a priori knowledge of antigen-specificity^27-32^. Most of the cell-based approaches were designed for antigen discovery of MHC class I-restricted CD8+ T cells, and their translation to MHC-II is not trivial. For instance, T-scan relies on secretion of GzmB by primary CD8+ T cells, and thus cannot be used for CD4+ T cells. The utility of cell-based, MHC/HLA-II, antigen discovery was demonstrated by Kisielow et al using their pMHC-TCR (MCR-TCR)^28^. The MCR-TCR complex allowed for the identification of cognate epitopes by iterative screening against libraries encoded through cDNA. Though sensitive, MCR-TCR method offers relatively limited throughput due to iterative screening.

With the increasingly widespread use of single cell RNA sequencing (scRNAseq) to interrogate T cell responses, it is paramount that T cell antigen discovery methods can be scaled to investigate 10s-100s of TCRs rapidly. Recently, several algorithms for computational antigen discovery have been reported, including GLIPH/GLIPH2, tcrdist/tcrdist3, and CoNGA^33-35^. These algorithms identify TCR specificity groups made up of TCRs that share sequence similarity and/or motifs, and are therefore predicted to share antigenic specificity. Therefore, if the antigenic specificity of a given TCR is known, its analogs can be identified and experimentally tested for binding to the same antigens. Recently, this approach, termed ‘reverse epitope discovery’ has been explored to leverage large datasets for comparison of TCR amino acid similarity^36^. Ultimately, Pogorelyy et al were able to identify public, immunodominant CD4^+^ T cell responses across 59 individuals. However, it remains challenging to predict the antigens of private clonotypes in private datasets. Furthermore, the method requires an existing dataset containing TCR sequences of known antigen specificity, highlighting the need for high throughput methods which synergize both experimental and computational approaches^10^.

Here we showcase a combination of several methodological advances in applying experimental and computational tools for antigen discovery. Firstly, we report a cell-based method for antigen discovery using SABR-IIs for mouse and human CD4+ T cells. Second, we show *de novo* identification of epitope specificities TCRs derived from scRNAseq data in a mouse model of Type 1 Diabetes. Finally, we demonstrate that experimental antigen discovery can be paired with *post hoc* amplification by computational approaches. Together, we have developed an experimental and computational workflow to rapidly de-convolute scRNAseq-derived CD4+ T cells *de novo*. Our approach is broadly applicable, scalable, and reproducible, all towards the ultimate goal of comprehensively defining the CD4+ T cell repertoire.

## RESULTS

### MHC-II signaling and antigen-presenting bifunctional receptors

We have previously described SABRs, which are chimeric receptors containing an extracellular pMHC complex attached to an intracellular CD28-CD3ζ signaling domain. We demonstrated that SABRs can read out TCR-pMHC interactions, allowing the construction of SABR libraries for antigen discovery for class I HLA alleles^27^. Here, we created SABRs to present epitopes in MHC-II alleles, by covalently linking the epitope to the β-chain of MHC-II that is attached to the CD28-CD3ζ signaling domains downstream, along with a 2A peptide-linked MHC-II α-chain (**Fig 1A, B**). To test if SABR-IIs can present epitopes to TCRs and induce a signal, we expressed them in NFAT-GFP-Jurkat cells (a kind gift from Dr. Yvonne Chen and Arthur Weiss) using lentiviral vectors. We constructed murine SABR-IIs presenting epitopes in I-Ab, I-Ad, and I-Ag7 (Ova, ISQAVHAAHAEINEAGR^37^; ATEG, ATEGRVRVNSAYQDK^38^; 2.5mimo, YVRPLWVRME^39^ respectively) (**Fig S1A**). NFAT-GFP-Jurkat cells express GFP upon NFAT activation and translocation downstream of CD3ζ activation. We co-incubated the SABR-II-expressing NFAT-GFP-Jurkat cells with a separate population of Jurkat cells expressing either the Bdc2.5 TCR (recognizes I-Ag7-2.5mimo), OT-II TCR (recognizes I-Ab-Ova), 5-4-E8 TCR (recognizes I-Ad-ATEG), or no TCR. Robust GFP and CD69 expression in SABR-II-expressing NFAT-GFP-Jurkat cells was observed 18-20h later only in the correctly paired assays. (**Fig 1C and Fig S1B**). Building upon this, we asked if SABR-IIs could be used to present a library of epitopes for CD4^+^ T-cell antigen discovery. To that end, we constructed a SABR-II library to present islet-derived epitopes in I-Ag7 by curating a list of 4,075 published epitopes from the Immune Epitope Database (IEDB, iedb.org)^40^ and a study by Wan et al^41^(**SUPP FILE 1**). Of note, this defined library consisted of unmodified epitopes derived from endogenous proteins, synthetic mimotopes, deamidated epitopes, as well as Hybrid Insulin Peptides (HIPs) that exist only as a posttranslational modification and are not genetically encoded *in vivo*^*42,43*^. The epitope library was inserted into the I-Ag7-SABR-II backbone through pooled oligonucleotide synthesis, amplification, and ligation-free cloning (**Fig 1D**). The I-Ag7-SABR-II-library was then expressed in NFAT-GFP-Jurkat cells. As a proof-of-concept, we performed co-incubation assays with Jurkat cells expressing the BDC2.5 TCR and sorted GFP+CD69+ cells (**Fig S2A**). We extracted the genomic DNA from sorted cells, amplified the SABR portion of the integrated proviruses, and subjected the amplicons to Illumina sequencing (**Fig 1D, S2B**). Sequence reads were aligned to the I-Ag7-SABR-II backbone and the corresponding epitopes were scored based on their reads. For each TCR under investigation, two metrics were used to determine epitope enrichment (**Fig S2C**): RANK, the average difference between the relative ranks of each epitope in the library for a given TCR against an unsorted library; and FRACTION-READS, the average difference between the contribution of each epitope to the overall library abundance for a given TCR against a mock-sorted library. Triplicate assays were set up for each TCR and used to determine the average RANK and FRACTION-READS. Relative enrichment of each epitope was determined by subtracting the RANK and FRACTION-READS obtained from the mock sort from the experimental TCR sort (**Fig S2C**). Screening the Bdc2.5 TCR against the I-Ag7-SABR-II-library resulted in an enrichment of epitopes containing the WXRM(D/E) motif (**Fig 1E, S3A, B**), a well characterized trait of the Bdc2.5 TCR^44^, along with several known cognate epitopes of the Bdc2.5 TCR^39,45,46^ (**Fig S3A, B)**. Using SABR co-incubation assays described above, we confirmed several epitopes identified by us are indeed recognized by the BDC2.5 TCR. To determine the sensitivity and specificity of SABR screens, we used a set of 10 known cognate epitopes and 10 known irrelevant epitopes. Combined analysis of the rank enrichment of epitopes across 7 independent screens demonstrated the ability of I-Ag7-SABR-II-library to both rigorously and robustly identify cognate epitopes in SABR-II screens (**Fig 1F & G**). Together, these results demonstrate the ability of SABR-IIs, to successfully read out pMHC-II-TCR interactions and serve as a method for CD4^+^ TCR antigen discovery.

**Fig. 1.**
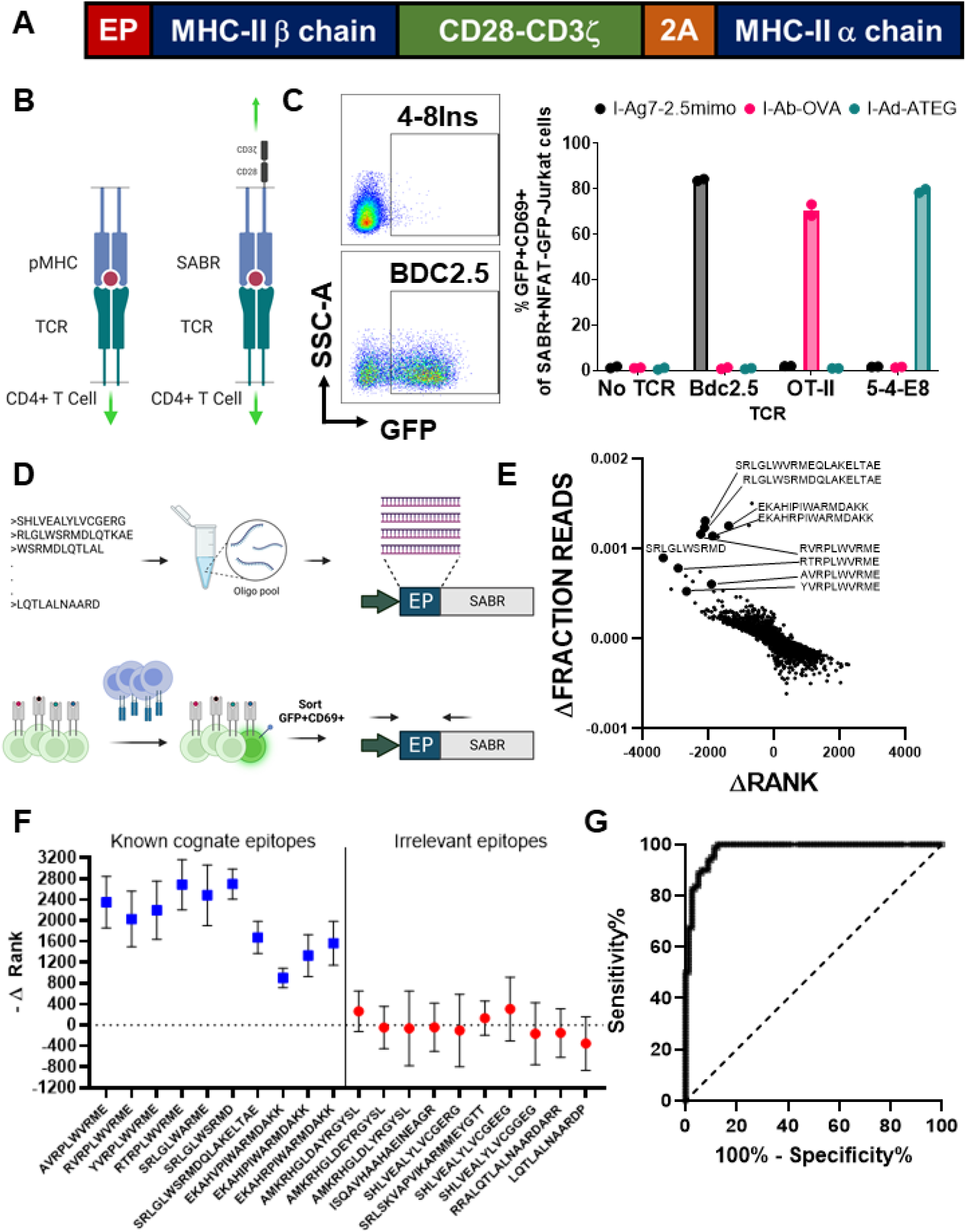
SABR-IIs for identifying cognate TCR-pMHC interactions and antigen discovery. **A**. A schematic of SABR-II constructs. **B**. Signaling directionality between both a physiological pMHC:TCR (left) and a SABR-II:TCR (right) with cognate interaction. **C**. Representative and summary plots for GFP and/or CD69 expression from SABR-expressing NFAT-GFP-Jurkat cells after culture with TCR-expressing Jurkat cells. Bar graph plots mean and s.d. for 2 technical replicates. **D**. Schematic overview of pooled oligonucleotide cloning for SABR-II libraries (top) and co-culture assays for SABR-II screens. **E**. Representative plot from I-Ag7-SABR-II library screen of the Bdc2.5 TCR. Each dot in the plot is an epitope from the library. X-axis represents the mean Delta-FRACTION-READS and Y-axis represents the mean Delta-RANK for each epitope. Several representative epitopes are indicated with larger dots and the epitope sequences are listed. **F**. Accuracy and specificity of SABR-II libraries. The Delta-RANK for known (blue boxes) and irrelevant (red circles) epitopes across 8 independent I-Ag7-SABR-II library screens of the Bdc2.5 TCR are shown. Mean and s.d. for are plotted. **G**. Receiver Operating Characteristic curve corresponding to the 8 independent I-Ag7-SABR-II library screens of the Bdc2.5 TCR.

### Single cell profiling of islet infiltrating CD4^+^ T cells

Having constructed and validated the I-Ag7-SABR-II-library, we reasoned that a pool of private CD4^+^ TCRs could be interrogated for their antigen specificity *de novo*. To that end, we used the NOD mouse model which recapitulates many features of T1D and shares several autoantigens with T1D patients^43,47,48^. Despite several known antigens, the overall antigenic landscape of islet infiltrating CD4+ T cells in NOD mice remains undefined. A recent study mapped the immune infiltrates in the islets of NOD mice by scRNAseq, but did not map their TCR repertoires, precluding antigen discovery^49^. Building upon this, we performed scRNASeq with V(D)J enrichment on T cells from individual pancreatic islets of 6-, 8-, and 10-week-old NOD mice. We sorted Thy1.2+TCRβ+ T cells from 3-4 mice at each time point, combined them using TotalSeq cell hashing oligonucleotides, and proceeded to scRNAseq using 10x Genomics platform. In total, T cells from 11 mice were sequenced in 3 batches and the data were pooled for analysis (**Fig S2A**). Hierarchical clustering in Seurat^50,51^, followed by bioinformatic gating on CD4+ T cells and re-clustering, revealed 7 distinct CD4+ T cell clusters with no obvious bias between mice (**Fig 2A and S4B**). Next, we integrated TCR clonotypes with the transcriptomes using scRepertoire^52^, and identified the clonally expanded populations of CD4^+^ T cells (**Fig 2B**). Clonal expansion was categorized as single (1 clone per TCR), low (2-9 clones per TCR), or medium (>10 clones per TCR). Clonal expansion was evident in clusters 0, 3, 4, 5, & 6, each demonstrating its own unique pattern of differentially expressed genes (**Fig 2C**). Generally, clonal expansion correlated with the expression of activation and exhaustion markers (Nkg7, Ccl5, Lag3, and Tigit), whereas naïve T cell markers (Sell, Ccr7) coincided with un-expanded populations. For the purpose of this study, we used TCR clonality to identify the top expanded TCRs. We reasoned that clonally expanded cells within the islets were the most likely to target islet antigens and contribute to β-cell destruction. Therefore, we focused on clonal expansion as the sole criterion for selecting TCRs for antigen discovery. Interestingly, expanded clones did not segregate solely based on their gene expression as indicated by the high degree of clonal sharing between CD4^+^ TCR clusters determined by the Morisita-Horn Index (**Fig 2D**). Clonally expanded TCRs showed increased expression of Lag3, corresponding to their restrained phenotype that was described previously^53^. Further investigations into the transcriptional signatures of expanded T cells are reported in an accompanying manuscript (Xiao, Rohimikollu, and Rosengart et al). Specifically, we identified the top 40 clonally expanded TCRs, corresponding to 20 from each of the 8- and 10-week cohorts of NOD mice (**SUPP FILE 2**). These TCRs were derived from 5 mice, and showed a slight skew towards certain Vα and Vβ alleles, as has been described previously. We then reconstructed the TCRs using a homebrewed python script that reconstructs full TCRα/β chains using the IMGT TCR allele dataset (See Methods)^54^. The reconstructed TCR genes were synthesized through commercial vendors and sub-cloned (**Fig 2E**) into the pMIG-II-IRES-GFP vector containing a partial Cβ chain derived from the BDC2.5 TCR. TCRs in the pMIG-II vector were then packaged intro retroviruses and expressed in Jurkat cells. Surface expression was confirmed by staining for murine TCRβ followed by flow cytometry. For TCRs with low transduction levels, we enriched the TCRβ+ population using either FACS or magnetic selection (**Fig 2E**). Altogether, scRNAseq of islet infiltrating T cells in NOD mice revealed clonally expanded populations that yielded TCRs for antigen discovery.

**Fig 2.**
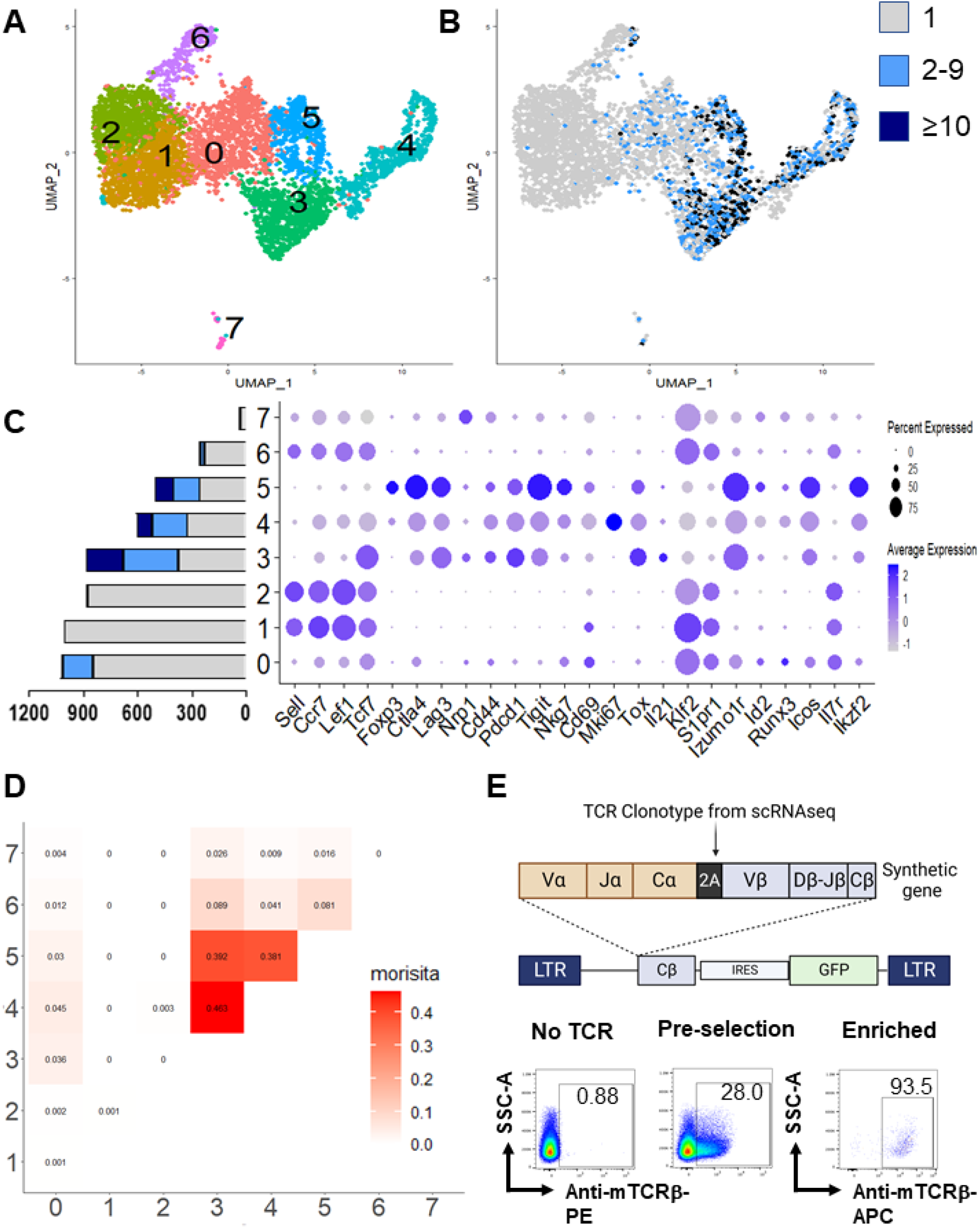
Single-cell RNA-sequencing of islet infiltrating CD4+ T cells. **A**. UMAP representations of islet infiltrating CD4+ T cells from 6-, 8-, and 10-week old NOD mice. Hierarchical clusters generated by Seurat are shown in different colors and numbered according to the cluster. **B**. Overlay of clonal expansion on the clusters. Grey dots represent cells with unique clonotypes, Light blue dots represent low (2-9 clonotypes) expansion, Dark blue dots represent high (>10 clonotypes) expansion. **C**. Cluster composition (left bar graph depicts cell number on x-axis, colors denote clone size from panel B) and differential gene expression map across clusters. Individual genes are represented on X-axis and scaled according to the average expression. **D**. Morisita-Horn index plot comparing TCR sequences across each cluster. **E**. Schematic (top) of TCR cloning strategy into pMIG-II backbone along with representative flow cytometry plots (bottom) of murine TCR levels before and after enrichment.

### SABR-II libraries identify cognate epitopes of CD4^+^ TCRs *de novo*

We proceeded with a systematic screening of the cloned TCRs against the I-Ag7-SABR-II-library in the manner described above (**Fig S5A and B**). High confidence hits for 10 of the screened TCRs across 3 separate antigen specificities were identified de novo (**Fig 3A**). These antigens were: two HIPs InsulinC-ChgA (LQTLALQSRMD) and InsulinC-IAPP (LQTLALNAARDP), and InsulinB9:23 (SHLVEALYLVCGERG). We validated the reactivity by generating SABR-IIs presenting solo epitopes corresponding to the putative hits and confirming their recognition by the corresponding TCR as described above (**Fig 3B and S6A**). All the putative hits identified in the screen were validated, reaffirming the robustness of the SABR-II library to determine the ligands of MHC-II-restricted TCRs *de novo*. Further validations using in vitro mIL-2 secretion by TCR-expressing 5KC reporter cells^13^ or CD25 expression by TCR-expressing splenic CD4+ T cells upon stimulation with the cognate epitope were performed (**Fig S6B and C**). To our knowledge, this is the first report of cell-based epitope discovery of TCRs derived from scRNA seq data. Interestingly, visualization of the cells corresponding to each de-convoluted TCR clone did not reveal overt differences in the phenotype of cells recognizing the three different antigens (**Fig 3C**). Taken together, these results indicate that SABR-II libraries can successfully identify cognate epitopes of CD4^+^ TCRs amongst thousands of epitopes for TCR-directed antigen discovery starting simply from a TCR sequence and with little a priori knowledge of target antigens.

**Fig 3.**
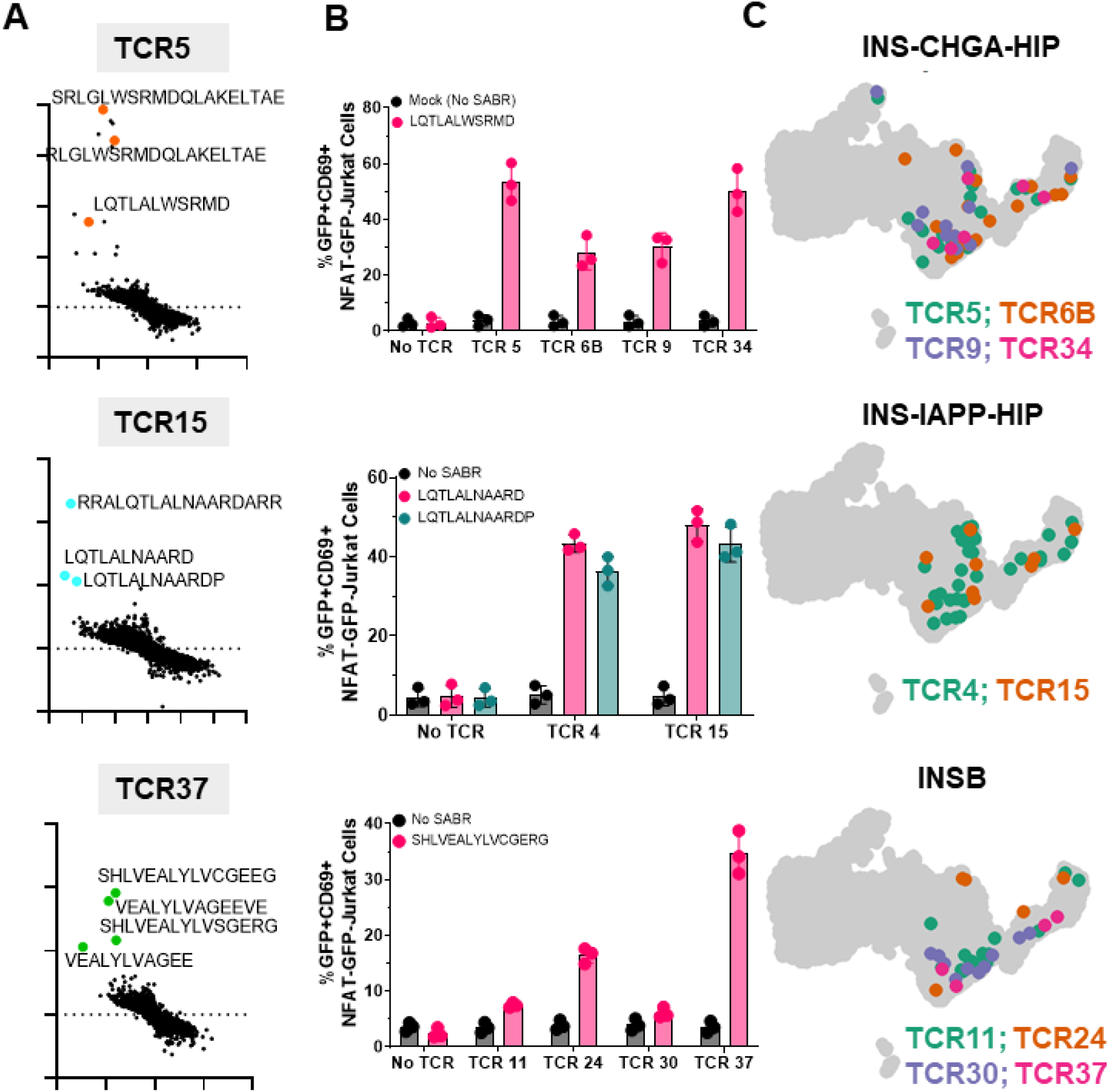
De novo identification of the cognate epitopes of expanded CD4+ T cells. **A**. Representative SABR-II screen results for three TCRs are shown. Putative hits are indicated with the epitope sequences and larger dots. **B**. Single SABR-II coincubation assays with TCR expressing Jurkats with NFAT-GFP-Jurkats expressing SABR-IIs presenting their putative hit epitope(s). GFP+CD69+ cells in co-incubation assays at 18 hours are quantified. Bars show mean and s.d. from 3 biologically independent experiments. **C**. Projection of antigen-specific CD4+ T cell clones onto the Seurat UMAP of islet infiltrating CD4+ T cells. Indicated TCRs and their cognate epitopes are denoted.

### *In silico* TCR similarity predictions amplify SABR-II antigen discovery

We hypothesized that computational grouping of TCR specificities may reveal closely related TCRs that potentially recognize the same epitope(s), similar to the reverse epitope discovery approach (**Fig 4A**). In absence of experimental antigen discovery, mere grouping of TCRs is not informative. However, armed with SABR-II-mediated de-convolution of the 10 TCRs from the scRNAseq dataset, we hypothesized that TCRs that co-cluster with these TCRs bind to the same antigens. To test this, we used 3 TCR similarity search algorithms: grouping of lymphocyte interactions by paratope hotspots (GLIPH2)^34,55^, distance measure on space of TCRs that permits clustering and visualization (tcrdist3)^33^, and clonotype neighbor graph analysis (CoNGA)^35^. All three algorithms take slightly different approaches to group TCR sequences and generate clusters of TCR sequences that share high sequence similarity. In addition, CoNGA considers the transcriptional similarities among T cell clones. These analyses identified 16 TCRs that co-cluster with 4 of the experimentally de-convoluted TCRs. These TCRs, referred to as ‘analogs’ hereon, were cloned and expressed as described above. We performed co-incubation assays using single SABRs and observed that 6/16 TCRs recognized the same epitopes as the parental TCRs. (**Fig 4B**). As a result, we were able to identify the cognate epitopes of 6 additional TCRs from our dataset which had otherwise not been selected for SABR-II screening based on our clonal expansion cutoff. Interestingly, the computationally identified and experimentally validated TCRs shared similar phenotypes as the experimentally de-convoluted TCRs (**Fig 4C**). Therefore, we demonstrated that computational TCR similarity determinations could amplify experimental antigen discovery, leading to a total of 16 TCRs de-convoluted de novo.

**Fig 4.**
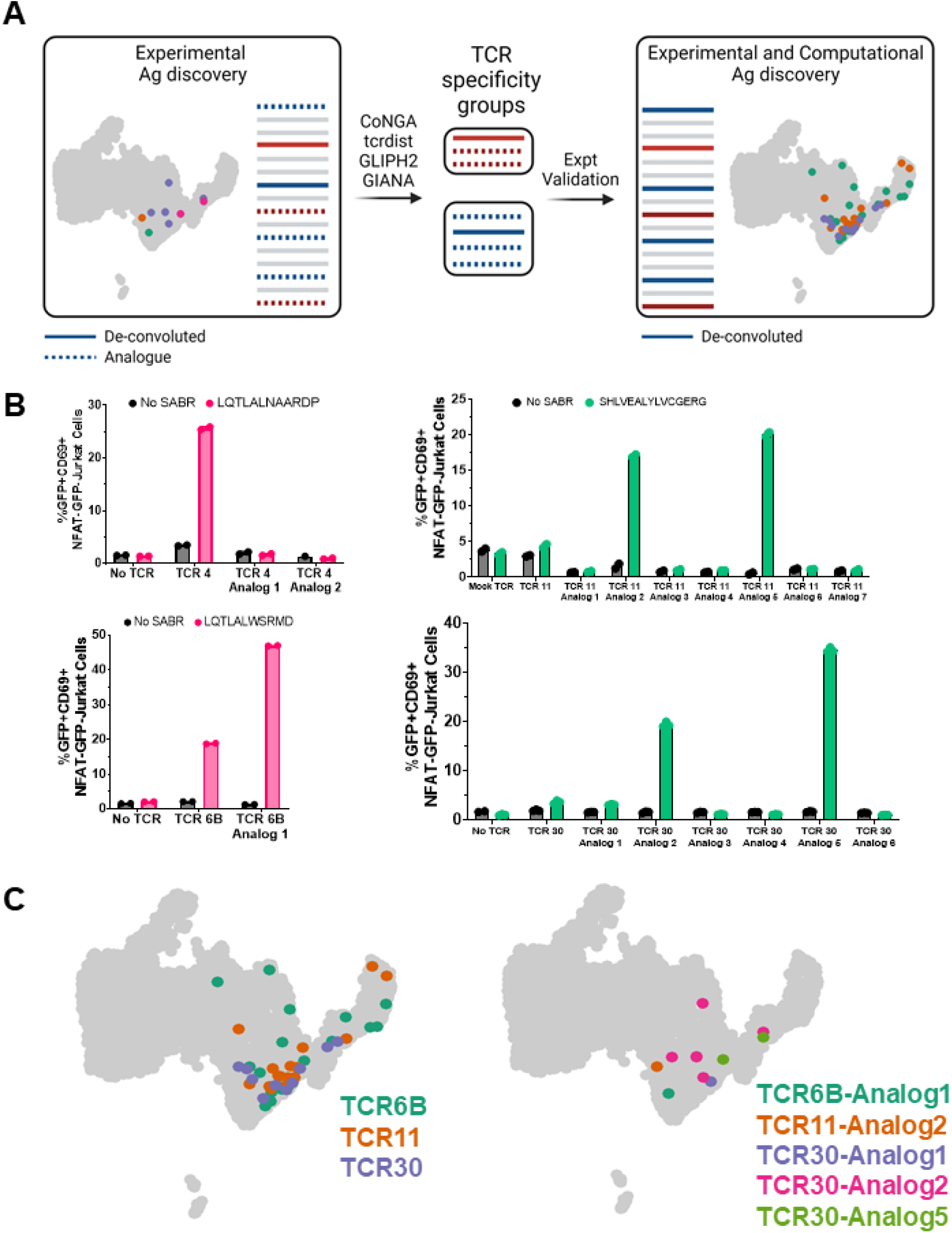
Computational prediction of antigen-specificity amplifies SABR-II antigen discovery. **A**. Schematic of the workflow for computational prediction of shared antigen specificity following experimental deconvolution. Solid red and blue lines indicate experimentally de-convoluted TCRs, dashed blue and red lines indicate potential analogs that may share specificity with the experimental TCRs, Grey lines indicate remaining TCRs. **B**. Single SABR assays similar to Fig 3B for four TCRs are shown here. Mean and s.d. of quantified GFP+CD69+ cells plotted from 2 technical replicates. **C**. Projection of antigen-specific CD4+ T cell clones onto the UMAP after validation of predicted antigen specificity (right) compared to original clones (left).

### HLA-SABR-IIs can be used to identify the cognate antigen(s) of human TCRs

The ability of SABR-IIs to identify cognate antigens for unknown murine CD4^+^ TCRs naturally led us to hypothesize that they could also be wielded for human antigen discovery. The ability to begin with only a TCR sequence and a SABR-II library allows for human samples to be maximized as they are often scarce. This is especially relevant for T cells that infiltrate the pancreata of human T1D patients which have only been collected from a relatively small number of donors^56^. To demonstrate humanized SABR-IIs, we generated SABR-IIs to present the InsB9:23 epitope (SHLVEALYLVCGERG) in HLA-DQ8 (DQA1*0301:DQB1*0302, an HLA-II allele that is associated with increased risk of T1D and Celiac disease^57,58^) to human TCRs. We confirmed the ability of the DQ8-InsB8:23 SABR-II to present the epitope to two previously described, T1D patient-derived TCRs GSE.6H9 and GSE.20D11^59^. As expected, high frequency of GFP+CD69+ cells was found only when the TCRs interacted with the InsB9:23 epitope, and not a control Hen Egg Lysozyme epitope. To test if HLA-DQ8-SABR-II could be used for antigen discovery, we curated a list of InsulinB, InsulinC, and HIP epitopes published by Wiles et al^60^ and cloned them into the DQ8-SABR-II backbone using the same cloning strategy as the I-Ag7-SABR-II-library (**SUPP FILE 3**). As a proof-of-concept, we screened both the GSE.6H9 and GSE.20.D11 TCRs expressed in Jurkat cells against the DQ8-SABR-II-library as described above. As expected, InsulinB9:23 was a top hit for the cognate epitope of both TCR GSE.6H9 and GSE.20.D11 (**Fig 5B**). Thus, we demonstrated that SABR-IIs can be used both for determining cognate TCR-pMHC interactions and screening large epitope libraries for de novo CD4^+^ T antigen discovery across both murine and human TCR repertoires.

**Fig 5.**
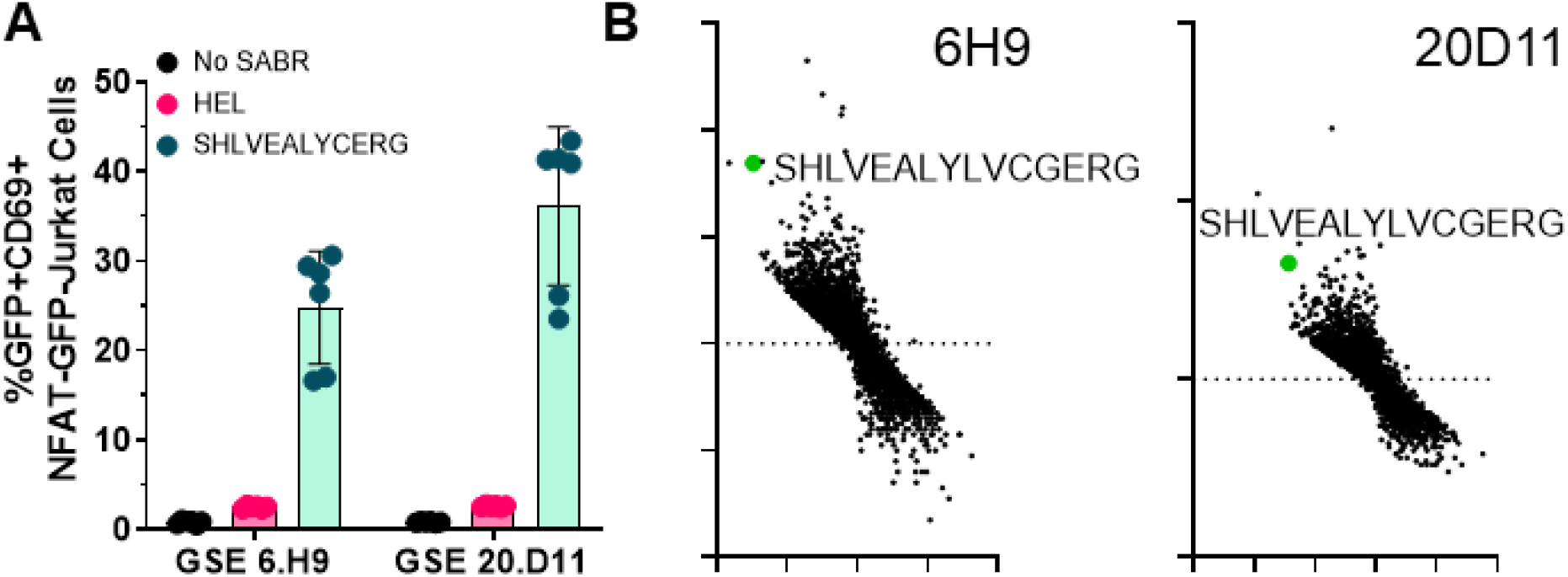
Proof of concept for HLA-SABR-IIs. **A**. Single SABR co-incubation assays of TCR expressing Jurkats with NFAT-GFP-Jurkat cells expressing InsB9:23 in HLA-DQ8 SABR-IIs. Mean and s.d. are plotted from 3 biological replicates. **B**. Representative plots from Jurkat cells expressing two known InsB9:23 reactive TCRs screened against the HLA-DQ8-SABR-II library. The InsB9:23 epitope enrichment is indicated by larger green dots.

## DISCUSSION

In this manuscript we present an integrated workflow to identify the epitopes of CD4+ T cells de novo. This workflow combines SABR-II libraries, TCRs reconstructed from scRNAseq data, and computational TCR grouping tools for antigen discovery. Using this workflow, we identified the cognate antigens of 16 islet-infiltrating TCRs from NOD mice de novo.

Our method has several advantages over current state-of-the-art methods. Firstly, SABR-II libraries are a robust method for encoding a large number (1000s-10,000s) epitopes for screening. In contrast to the MCR-TCR method, we can readily and precisely define epitope libraries and can screen TCRs in a single round of enrichment. While in this study, we use T cell lines to express SABRs, other non-T cell types can also be used, unlike MCR-TCRs, possibly expanding antigen discovery to professional APC based platforms. Importantly, SABR-II libraries have the ability to encode for several post-translational modifications, which is not possible through MCR-TCRs, as they derive epitopes from cDNA libraries. This is well demonstrated by the encoding for and identification of reactivity to HIPs, which have been recently identified as autoantigens in T1D but are generated by aberrant post-translational modifications^42,43^. To our best knowledge, this is the first report of a cell-based epitope discovery method that has encoded PTMs in addition to endogenous epitopes. The ability to start from and reconstitute TCRα/β sequences means that precious human samples are not wasted and can be assayed using additional methods. Furthermore, starting from scRNASeq has the built-in advantage of leveraging the transcriptional information for each clone of an identified specificity, not limited by a few phenotypic surface markers or agnostic of the T cell’s function altogether. This allows one to profile T cell specificities in an unbiased manner, unlike antigen-directed approaches that potentially over-represent T cells with greater function. Furthermore, this approach allows interrogation of specificities for T cells that are often functionally inert in vitro, such as regulatory T cells. While we have chosen to profile the top expanded T cell clones in this study, we envision that future efforts (such as those described in the accompanying manuscript by Xiao, Rohimikollu, and Rosengart et al) can be focused on specific phenotypes of interest, for instance, regulatory T cells. In this way, both the environment from which the T cells are sampled and the properties of the T cells themselves will help further shape hypothesis-driven antigen discovery. This will be especially important for further class-II antigen discovery in autoimmune diseases such as T1D and surveying the regulatory T cell compartment. The ability to amplify antigen discovery using related TCRs by leveraging existing computational methods not only validates their utility but generates a positive feedback loop for increased repertoire profiling and validation of TCR specificity. This will lead to an overall enlargement of the known epitope-specific TCR repertoire, and provide incorporation of orthogonally obtained datasets for de novo antigen discovery. Finally, SABR-IIs in conjunction with SABRs, allow parallel antigen discovery for CD4+ and CD8+ T cells with the same experimental setup, which is unprecedented.

We do wish to highlight the current limitations of our technique. As with the current cell-based epitope discovery methods, SABR-IIs cannot match the scale of yeast display which can reach up to 10^8^ epitopes for profiling. Therefore, while not required, certain a priori criteria such as MHC binding prediction, tissue expression patterns, or known immunopeptidomic datasets greatly enhance SABR-II library design. SABR-II screens are currently performed as “one against many” assays, accommodating a single TCR co-incubation against the library. With multiplexing, we can routinely screen 10s of TCRs in a single day. Another current limitation is surveying TCR reactivity against multiple MHC or HLA restrictions simultaneously. However, we envision that this could easily be overcome by combining NFAT-GFP Jurkat cells of the same epitope library in two or more alleles as the MHC sequences themselves could serve as the differential barcode. The computational prediction tools we used here also pose inherent limitations to our workflow. As shown, 10/16 computationally predicted TCRs did not recognize the same antigens as the parental TCRs. This may be due to the erroneous calling of clonotypes, or due to the analogs binding variations of the epitopes tested here. Either way, while we were able to amplify experimental antigen discovery, caution must be taken to not presume that prediction equals actual binding.

While we showed *de novo* identification of 10 top expanded TCRs out of 40, we did not identify the cognate epitopes of the remaining TCRs. This could be due to several reasons. Firstly, we used a published MHC elution dataset, which inherently has high specificity but low sensitivity for detecting MHC-II bound epitopes. Building new SABR-II libraries based on tissue-specific gene expression may benefit by casting a wider net in search of cognate epitopes. In addition, a hallmark of numerous autoreactive diseases is the reactivity to post-translationally modified epitopes^61,62^. While we were able to encode hybrid and deamidated epitopes in our SABR-II libraries, we are developing approaches to incorporate a wider range of chemical modifications. Finally, the antigen sensitivity of class I SABRs is inherently lower than those of TCRs. We expect that SABR-IIs may also have a similar limitation where very low-affinity antigens do not generate a strong SABR signal and remain below the limit of detection without further modification such as the introduction of a disulfide trap to stabilize the MHC and fix weak binding registers in place.

In summary, this study demonstrates that wielding SABR-IIs for TCR-directed antigen discovery and amplifying discovery with existing computational methods is a powerful combination for understanding CD4^+^ T cell specificities. By increasing the ability to survey the T cell repertoire we envision a more comprehensive catalogue of the T cell “reactome” in numerous disease contexts, including autoimmunity, tumor immunity, host-microbe interactions, transplantation immunity, and antiviral immunity.

## ACKNOWLEDGEMENTS

We thank A.R. Cillo, C. Workman, J. Bridge, P. Thomas, S. Schattgen, S. Chiou, J. Das, H. Xiao, and H. Singh for scientific discussions and advice on experimental and computational techniques. We thank M. T. Leonard, K. Ford, K. Rankin, S. Rathod, L. Kublo, S. Martinez, and A. Parikh for technical assistance in the experimental antigen discovery pipeline. We thank T. Tabib and R. Lafyatis at the Single Cell Core, The Unified Flow Cytometry Core, and the Division of Laboratory Animal Research for aiding with scRNAseq, flow cytometry, and animal husbandry. NFAT-GFP-Jurkat cells were a gift from A. Weiss (University of California, San Francisco, San Francisco, CA, USA) and Y. Chen (University of California, Los Angeles, Los Angeles, CA, USA). The pCCLc-MND-X backbone and pCMV-RD8.9 were gifts from D.B. Kohn (University of California, Los Angeles, Los Angeles, CA, USA). 5KC cells were a kind gift from M. Nakayama (Barbara Davis Center for Diabetes, University of Colorado Anschutz Medical Campus). This research was funded by NIH/NIDDK New Investigator Gateway Award (1R03DK127447-01), NIH/NIDDK/dkNET New Investigator in Bioinformatics Award, Pittsburgh Autoimmunity Center for Excellence in Rheumatology Innovative Discovery Award (all to A.V.J.).

## AUTHOR CONTRIBUTIONS

P.Z. designed and performed experiments, analyzed and interpreted the data, wrote the manuscript. S.G. designed and performed the single cell RNA sequencing experiments. J.T., E. Z-M, L. H., R. R., N. T., S. A., performed experiments and assisted with technical procedures in TCR cloning, SABR screens, and validations. M. A. and D. A. A. V. provided reagents, guidance and scientific discussions. A. V. J. conceptualized the study, designed and performed experiments, analyzed and interpreted the data, and wrote the manuscript.

## METHODS

### Reagents and oligonucleotide primers

Reagents and Oligonucleotide primers methods can be found in **Supplementary File 4**. The lists of epitopes in the SABR-II libraries can be found in **Supplementary Files 1, 2, and 3**.

### Cell lines and peptides

Jurkat cells (ATCC) were cultured in R10 (RPMI 1640 media (Corning) supplemented with 10% FBS (Gemini Bio) and 10U/mL penicillin–streptomycin (Corning)). NFAT-GFP-Jurkat cells were a kind gift from Arthur Weiss and Yvonne Chen, and were cultured in R10 supplemented with 2 mg/ml Geneticin (Corning). HEK293T cells (ATCC) were cultured in D10 (DMEM (Corning) supplemented with 10% FBS (Gemini Bio) and 10U/mL penicillin–streptomycin (Corning)). 5KC cells were a kind gift from Dr. Maki Nakayama, and were cultured in IMDM with 10% FBS (Gemini Bio) and penicillin–streptomycin. All cell culture was performed at 37 °C with 5% CO2 in a humid cell culture incubator. Primary CD4+ T cells were isolated from spleens for NOD mice using Stemcell murine CD4+ T cell positive selection kit (Stemcell Technologies Inc).

### Mice

NOD/ShiLtJ (Strain 001976, The Jackson Laboratory) mice were purchased at the age of 4 weeks. The mice were fed autoclaved rodent breeder diet (T. R. Last). Female mice were used for scRNAseq and validation assays. All animal work was performed under the IACUC protocols in the AAALAC certified animal facility at the University of Pittsburgh.

### Construction of SABRs

SABRs were designed by assembling the individual component sequences in Snapgene (DNAstar). HLA allele chains were downloaded from IMGT and MHC allele chains were downloaded from UniprotKB. SignalP-5.0^63^ was used to predict the signal sequence and truncate it. The signaling domains were derived from the previously published SABR constructs^27^. Beta-chain-Signaling-2A-Alpha-chain fragments were assembled and codon-optimized using IDT’s codon optimization tool. BsmBI sites were replaced without affecting the amino acid sequences and EcoRI sites were added at the ends. A 2kb Stuffer fragment was also synthesized according to the previously published sequences. ORFs were synthesized as gBlocks (IDT) and assembled using PCR (KOD mastermix, Milipore Sigma) using the following primers: 2kb-Insert-gBlock-F; 2kb-Insert-gBlock-R; BsmBI-Insert-Fwd; ClassII-Alpha-Rev. The assembled full-length inserts were gel purified (Takara), digested with EcoRI (NEB), ligated in EcoRI-digested pCCLc-MND-X (A kind gift from Dr. Donald B. Kohn) and transformed using NEB-5alpha cells (NEB). Inserts were verified using MND_Input_Verify_F and MND_Input_Verify_R. Once full-length backbones were cloned, they were used to clone individual epitopes. To insert epitopes, SABR vectors were digested with BsmBI along with Alkaline Phosphatase to excise the 2kb stuffer fragment. Two complementary oligonucleotides, SABR-epitope-F and SABR-epitope-R, were synthesized for each epitope. Oligonucleotides were annealed to each other, phosphorylated, and ligated into the BsmBI-digested backbone (T4 Ligase, NEB), and transformed in NEB-5alpha cells (NEB). For cloning SABR libraries, oligonucleotide pools containing overhangs (Oligonucleotide epitope primer) were synthesized via Twist Biosciences. The pool was amplified using ClassII-Oligo-Fwd and ClassII-Oligo-Rev, and cloned in BsmBI-digested backbone using Infusion HD cloning (Takara). Bacteria were plated on LB-agar containing 100ug/ml carbenicillin (Life Technologies), grown overnight, and single colonies were selected for verification by Sanger sequencing (Azenta). Successful clones were used to inoculate liquid culture for overnight growth followed by plasmid minipreps (Zyppy miniprep kit, Zymo). Pooled libraries were subjected to maxipreps (Nucleobond Maxiprep EF kit, Takara).

### Single-cell RNA-sequencing of islet infiltrating T cells and analysis

NOD mice were euthanized by CO2 asphyxiation and immediately dissected for pancreas perfusion. Pancreas perfusion was performed under a dissecting microscope. Pancreatic duct was clamped using surgical clamps and 3 ml of 600 U/ml Collagenase dissolved in HBSS was injected using a 30G needle. Perfused pancreata were harvested and incubated at 37°C for 30 min. After the incubation, HBSS with R10 was added to quench collagenase. After washing twice with HBSS+R10, the tissue was plated on a 10 cm plate, individual islets were picked using a micropipettor. Islets were then incubated in dissociation buffer, centrifuged, and resuspended in the staining mix (1:500 dilution of anti-Thy1.2-BV605 + 1:500 dilution of Live/Dead-APC-Cy7, and 1:100 dilution of cell hashing TotalSeq antibodies (Biolegend)). After staining, the cells were resuspended in PBS+0.04% BSA and sorted on BD FACS Aria III sorter. After sorting the cells, they were counted and processed for scRNAseq. Cells were processing using 10x 5’ single cell gene expression kit v3 in a Chromium controller according to the manufacturer’s protocols. V(D)J enrichment was done using the single cell 5’ VDJ enrichment kit according to the manufacturer’s protocols. Libraries were sequenced on HiSeq4000 (Novogene Inc) with a 70:20:10 mix for gene expression:VDJ:hashing libraries. Sequence data were downloaded on the Joglekar laboratory server and aligned to the mouse genome (Mm10) using *cellranger* (10x Genomics Inc). TCR annotation was performed using *cellranger vdj* using mouse GRCm38 assembly. All three timepoints were sequenced and processed separately. Cellranger and cellranger vdj output files were used as inputs in Seurat^50,51^ for normalization, scaling, and dimensionality reduction. The packaged scRepertoire was used for TCR clonotype calling and analyses. The data were normalized using NormalizeData and scaled using ScaleData functions in Seurat. The scRepertoire^52^ functions combineTCR and combineExpression were used to add TCR clonotypes to each cell. HTODemux function in Seurat was used to demultiplex cell hashes and assign the correct mouse identity to each cell. At this point, all three timepoints were merged in Seurat using the merge function. After merging, integration was done using FindIntegrationAnchors and IntegrateData functions. Principle component analysis was performed using RunPCA. Top 20 principle components were used for Uniform Manifold Approximation and Projection, followed by cluster identification using FindNeighbors and FindClusters. CD4+ T cells were subsetted using FeatureScatter and CellSelector functions, and reclustered. Cluster markers were defined by FindAllMarkers function. Clonotype data were sorted according to expansion and exported as a csv file. UMAP representations with clonotypes were generated using highlightClonotypes function in scRepertoire. Differentially expressed genes were identified using FindMarkers function using DESeq2 statistics and represented using EnhancedVolcano function. For the accompanying manuscript (Xiao, Rohimikollu, and Rosengart et al), single (1), low (2-9), and medium (≥10) clonotypes were subsetted in Seurat and exported as Seurat objects for further analyses.

### TCR reconstruction and synthesis

TCR Vα, Jα, Vβ, and Jβ alleles along with CDR3α and CDR3β sequences were used as the input to reconstruct full length TCR sequences using the TCRgen_mouse.opt_v2.py script (available on Github). Mouse reference sequences were downloaded from IMGT. Full length TCR sequences (TCRα-2A-TCRβ) flanked by EcoRI site and truncated at the BlpI site in Cb were synthesized as gene fragments via Twist Biosciences. TCRs gene fragments were amplified using TCR-gene-Fwd and TCR-gene-Rev, and subcloned using pMIG-II vector containing the BDC2.5 TCR (Vignali laboratory) using EcoRI-BlpI. Successful cloning was verified using Sanger sequencing (Azenta).

### TCR similarity determinations

Exported clonotypes were used as inputs for GLIPH2^55^. For CoNGA, the merged dataset was exported as a .h5ad file and used as an input along with the cellranger vdj output file. CoNGA analysis was performed using default parameters^35^. Pairwise relative distances among TCRs were calculated using tcrdist3^33^. CoNGA, tcrdist3, GLIPH2 output files were searched manually for analogs that co-cluster with experimentally deconvoluted TCRs. Analogs were synthesized and cloned as described above.

### Generation and cloning of SABR libraries

To generate the I-Ag7 restricted SABR library, we combined all Immune Epitope Database epitopes with a published immunopeptidome generated by Wan et al^41^. Sequences were filtered remove all posttranslational modifications except deamidation and HIPs, and trimmed between 9-25 amino acid lengths. For the HLA-DQ8 library, non-contiguous epitopes from Wiles et al as well as all IEDB epitopes were combined to generate the epitope list. Epitope sequences were backtranslated using the backtranslate_fast.py script.

### Lentiviral vector production and transduction

Lentiviral vectors to express SABRs or TCRs were packaged via previously described procedures^27,64^. Briefly, a mixture of the lentiviral shuttle plasmid, pMDG-VSVG, and pCMV-RD8.9 (a kind gift from Donald B. Kohn) were transfected into HEK293T cells using TransIT-293 (Mirus Bio) and OPTI-MEM (Life Technologies). After 3 days, viral supernatant was collected and filtered through 0.45-μm syringe filters (Millipore). When possible, the freshly filtered virus was used to transduce 1×10^6^ Jurkat cells per ml of the virus. Occasionally, the virus was stored at –80 °C until use. For NFAT-GFP-Jurkat cells, Geneticin was added 24hr following transduction.

### Retroviral vector production and transduction

Retroviral vectors (pMIG-II) to express TCRs were packaged via previously described procedures. Briefly, a mixture of the retroviral shuttle plasmid, pRD114, and pHIT60 were transfected into HEK293T cells using TransIT-293 (Mirus Bio) and OPTI-MEM (Life Technologies). The following day, viral supernatant was collected and filtered through 0.45-μm syringe filters (Millipore). Transduction of 2.5×10^5^ Jurkat cells was performed using Retronectin (Takara) binding according to the manufacturer’s protocol.

### Coculture assays

For SABR screens, 5×10^5^ SABR expressing NFAT-GFP-Jurkat cells were labeled with cell trace violet (BioLegend) according to the manufacturer’s protocol before incubation with 5×10^5^ TCR expressing Jurkat cells in a round bottom 96-well plate for 16-20 hr. Cells were stained with anti-CD69-APC-Cy7 (Biolegend) and acquired on the Attune NxT flow cytometer (Thermo Fisher Scientific).

### High-throughput sequencing and analysis

Genomic DNA was extracted from sorted cells immediately after sorting, using the PureLink genomic DNA extraction kit (Life Technologies). The integrated SABR vectors were amplified with KOD polymerase (Millipore) and two rounds of amplification. In the first round, IDT-UD-SABR-C2-F and IDT-UD-SABR-C2-R were used to amplify the epitope. In the second round, UDI0001-R and UDI0001-F (representative of Index #1) were used to add Illumina Unique Dual Indexes (UDIs) to the amplicons. A different UDI was used for each sample. The reactions were pooled and purified with the Nucleospin gel and PCR purification kit (Takara). The purified PCR product was analyzed 2% agarose gel and subjected to sequencing on a HiSeq4000 (Fulgent Genetics). Unaligned reads generated by the sequencer were stored in FASTQ files. FastQ files were concatenated to generate one file for read1 and read2 each. The sequences were demultiplexed into individual indexes using demultiplex_dual.py. Epitopes were extracted and scored using epitope_extract_fastq_v1.1.py and merge_counts_split_v2.1.py. Delta Rank and Delta Reads were counted in Microsoft Excel.

### Statistical analysis

Flow cytometry plots were analyzed with FlowJo (Treestar). Statistical analyses and graphical representations were generated by Microsoft Excel (Microsoft) and GraphPad Prism (GraphPad).

### Reporting Summary

Further information on research design is available in the Nature Research Reporting Summary linked to this article.

### Data availability

All necessary scripts are deposited to Github (https://github.com/joglekar-lab/SABR-II), the sequencing data will be deposited to SRA. Plasmids and libraries will be deposited on Addgene. TCRs will be made available upon request, given the large number of them. Epitope specificities will be added to IEDB.

### Ethics statement

All animal work was performed as per institutional IACUC guidelines under an approved IACUC protocol (protocol # 20037102). All experimental work was performed according to the institutional IBC protocols.

---

**Fig S1.**
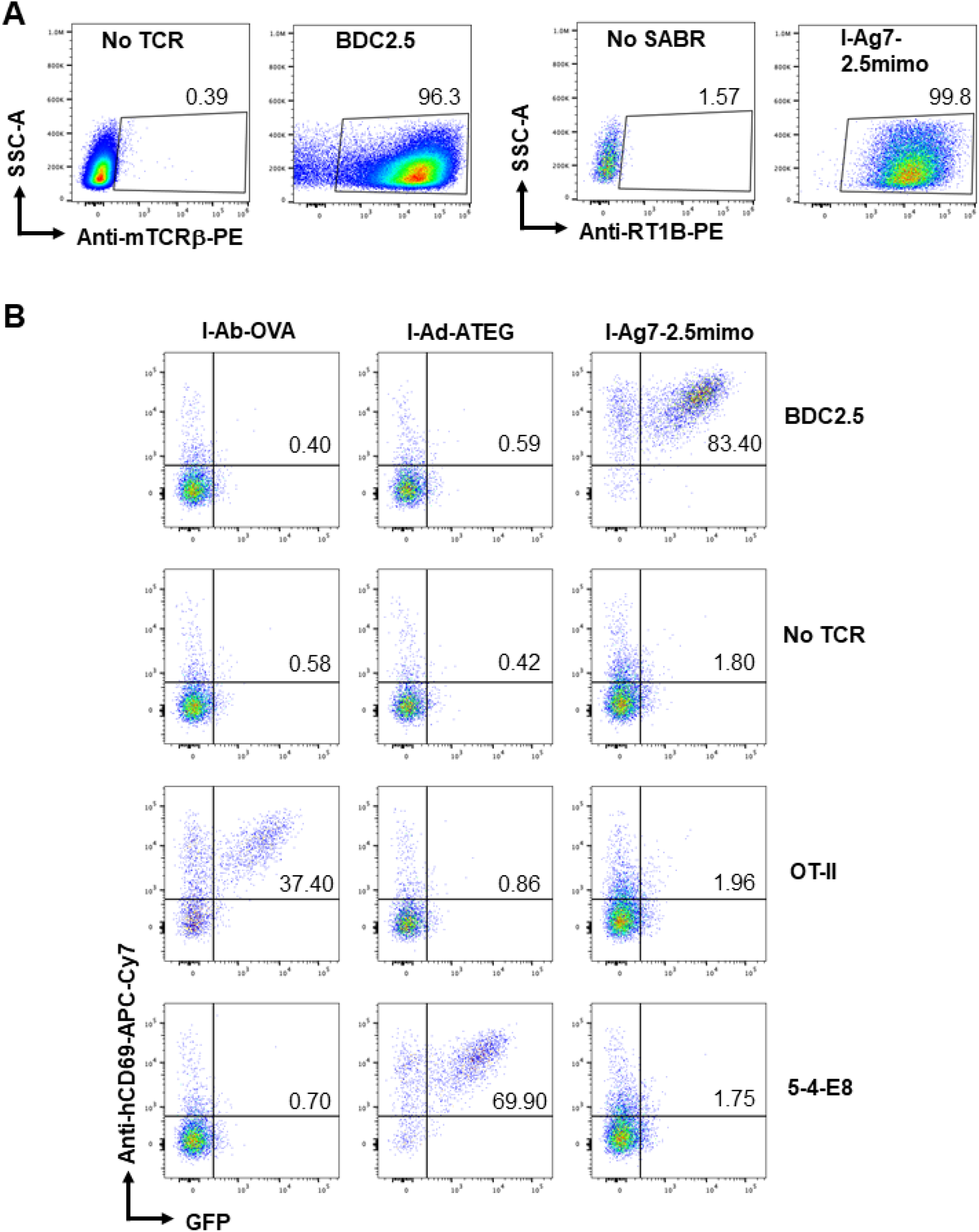
TCR and SABR construct expression and representative coincubation flow plots. **A**. Representative expression of murine TCRβ (Bdc2.5 TCR) and I-Ag7-2.5mimo SABR-II after transduction of Jurkat and NFAT-GFP-Jurkat cells respectively. **B**. Representative flow cytometry plots of SABR-II expressing NFAT-GFP Jurkat cells after co-incubation with TCR expressing Jurkats. The respective TCRs and SABRs are indicated.

**Fig S2.**
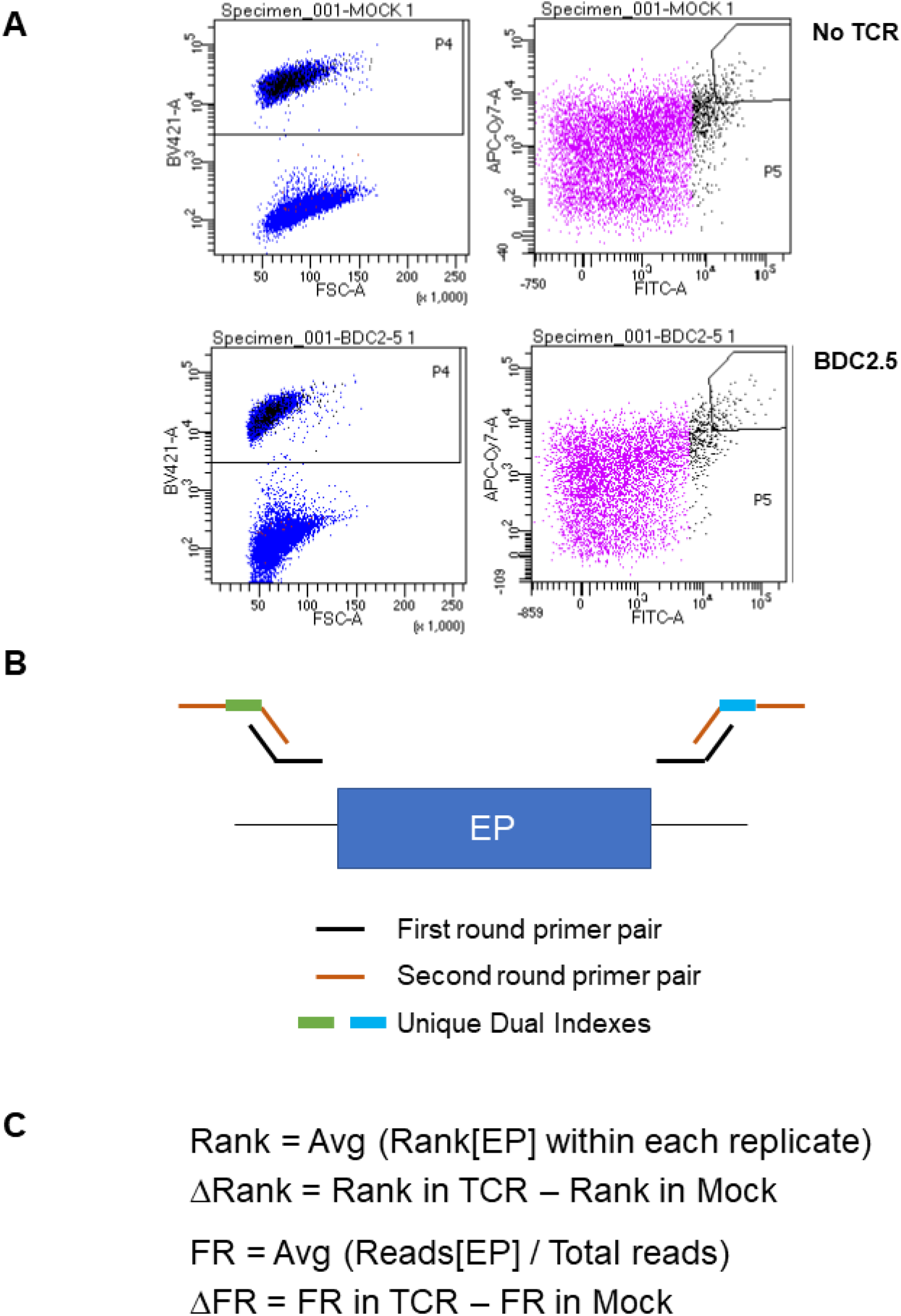
SABR-II library sort gating strategy and library indexing. **A**. Representative flow cytometry plots for SABR-II screen sorts. NFAT-GFP-Jurkat cells expressing the SABR-II library were labeled with cell trace violet, gated, and subsequently used to select top ∼2% of GFP/CD69 double positive cells for sorting. **B**. Schematics of the PCR strategy used for targeted reamplification of gDNA from sorted SABR-II library cells. **C**. Calculation of Δ-Rank and Δ-Fraction Reads (ΔFR) for epitopes represented in SABR-II library screens.

**Fig S3.**
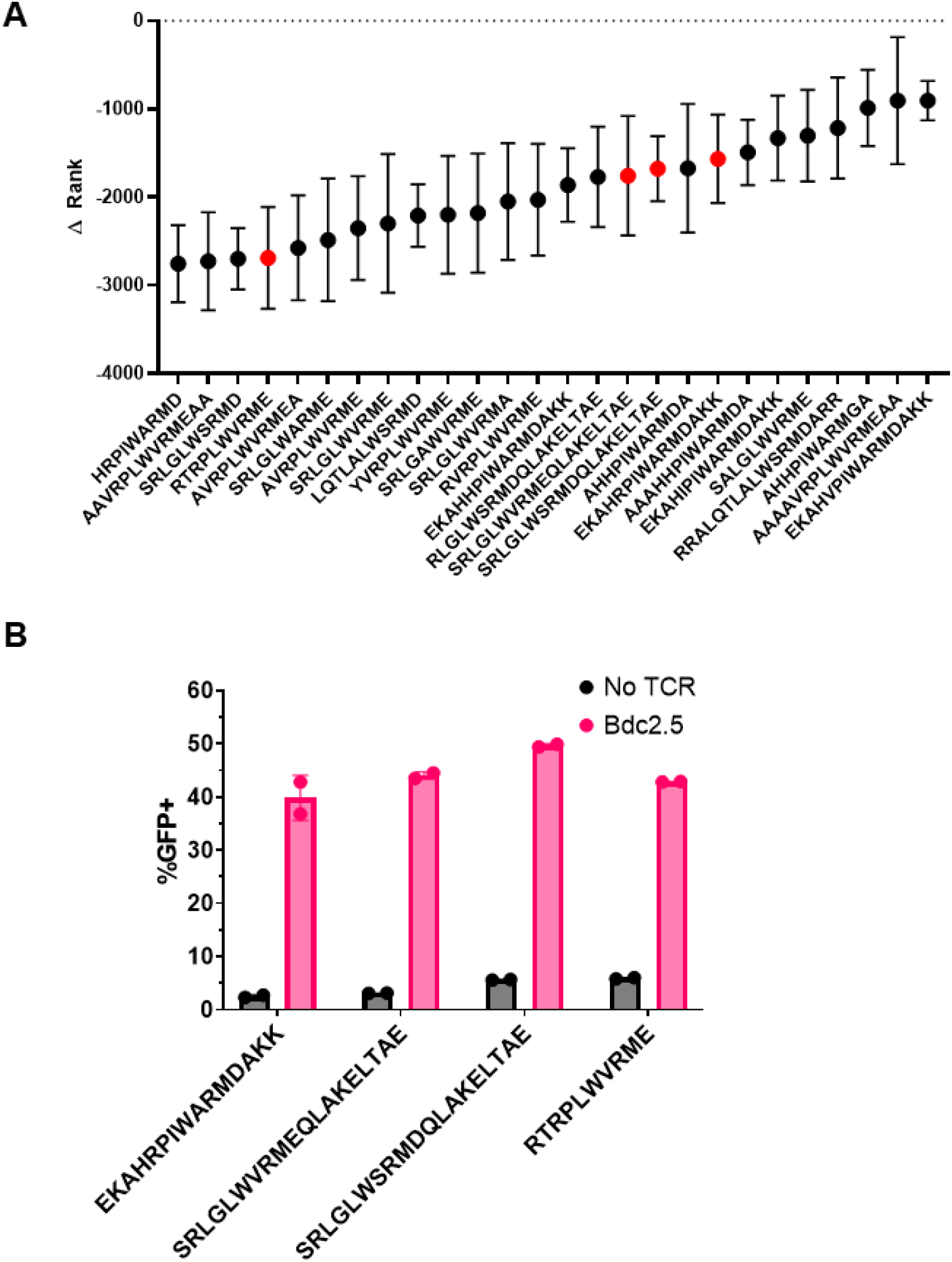
Top hits from multiple Bdc2.5 TCR I-Ag7-SABR-II library screens. **A**. Top 26 ranked putative epitope hits for the Bdc2.5 TCR across 8-independent I-Ag7-SABR-II library screens. Mean and s.d. are plotted for each epitope. Red dots indicate epitopes validated using single-SABR assays. **B**. Frequency of GFP+ NFAT-GFP-Jurkat cells expressing the putative hit epitopes in I-Ag7-SABR-IIs after 18 hr co-incubation with Bdc2.5 TCR expressing Jurkat cells. Bars show mean and s.d. from two technical replicates.

**Fig S4.**
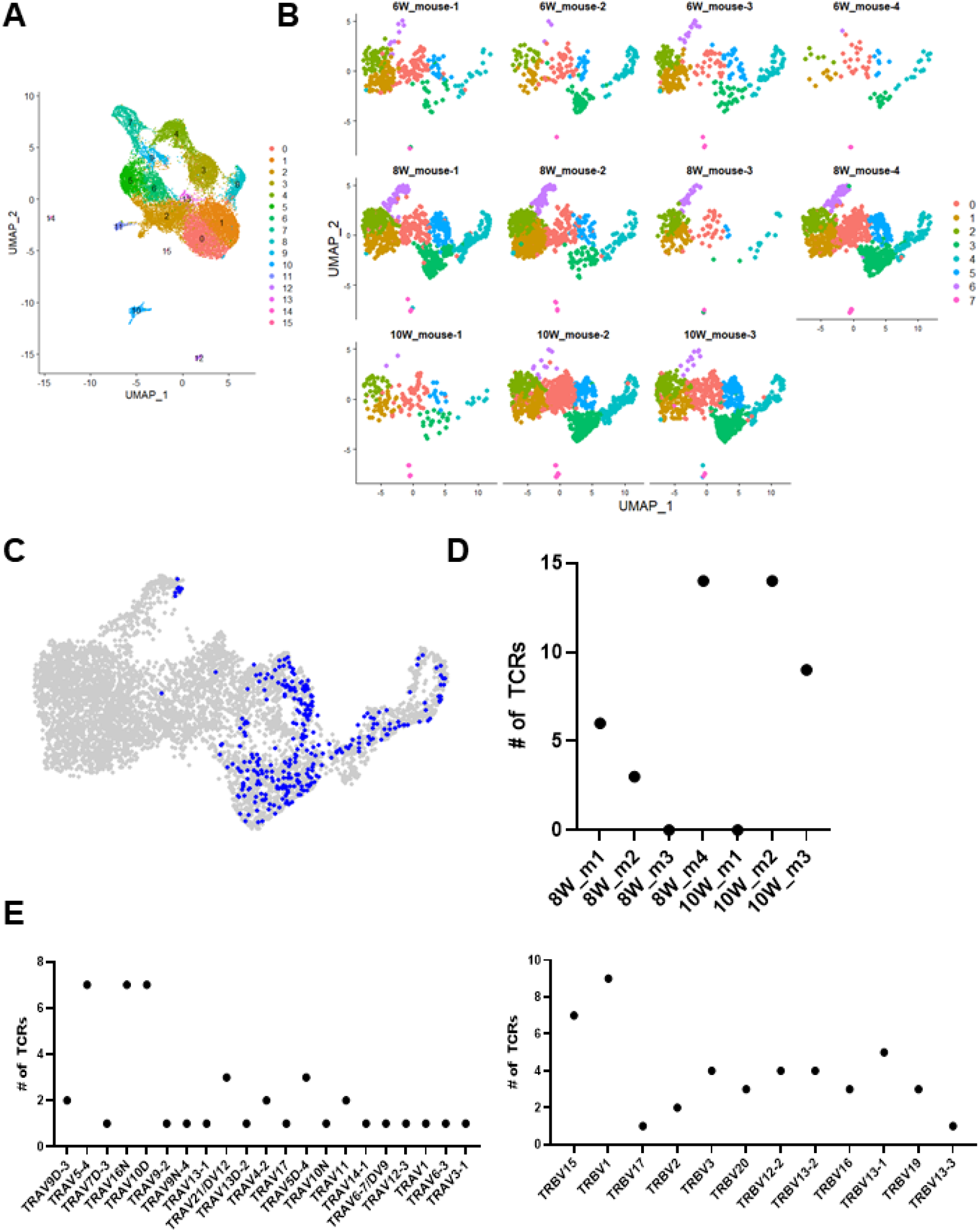
Single-cell RNA-Sequencing of islet-infiltrating CD4+ T cells from NOD mice. **A** Hierarchical clustering of total T cells across 11 mice from 6-, 8-, and 10-week old time points. **B**. Hierarchical clustering of CD4+ T cells from individual mice across 6-, 8-, and 10-week time points. **C**. Projection of top 40 expanded CD4+ T cell clones from 8-, and 10-week old NOD mice onto Seurat clusters. **D**. Distribution of top 40 expanded CD4+ TCR sequences across all mice. **E**. TRAV and TRBV usage from top 40 expanded CD4+ TCR sequences across all mice.

**Fig S5.**
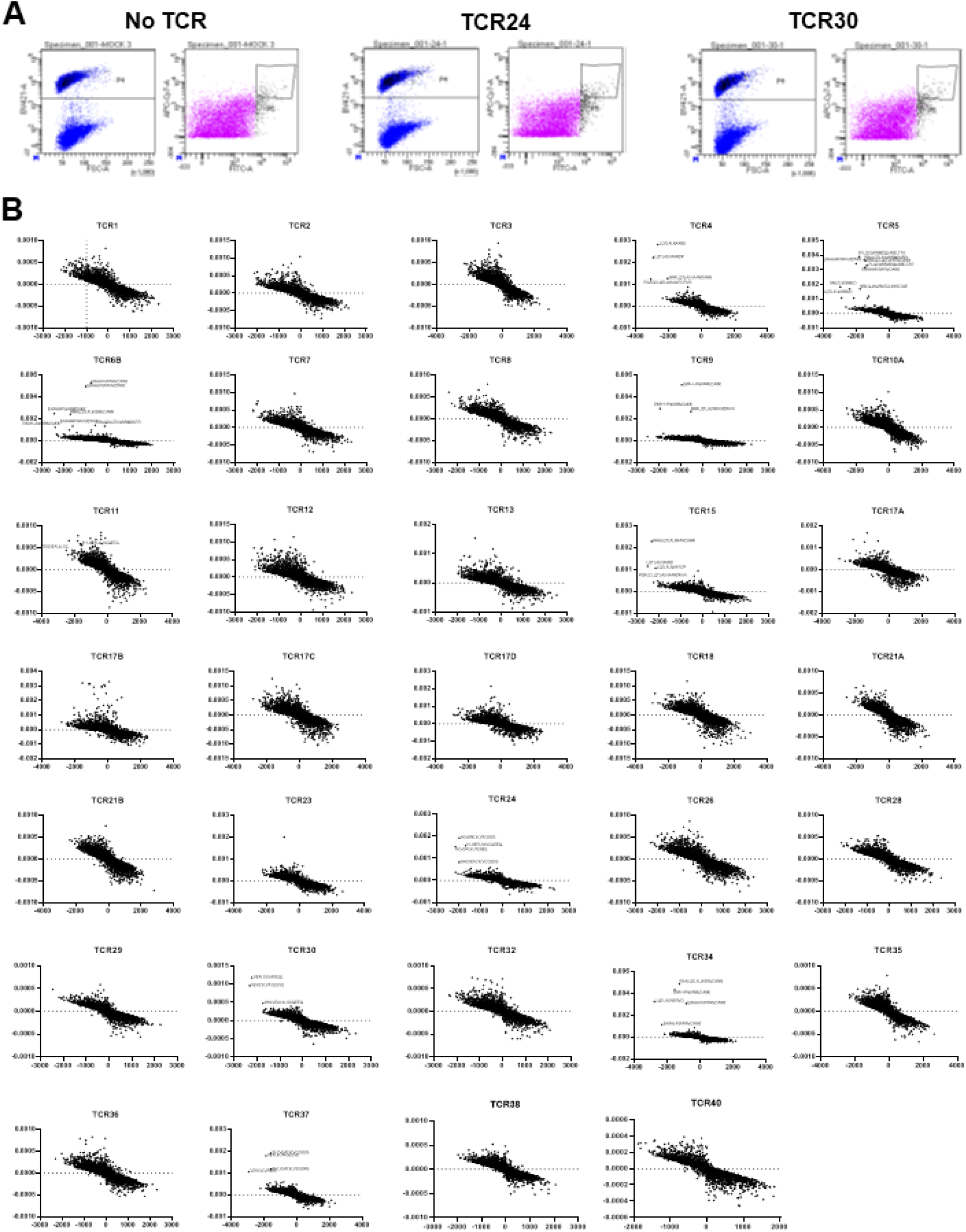
All I-Ag7-SABR-II library screen results. **A**. Representative flow sort gating for cell trace violet labeled NFAT-GFP-Jurkat cells expressing the I-Ag7-SABR-II library after co-incubation with Jurkat cells expressing TCRs. Top 2% of cells expressing GFP and CD69 were sorted. **B**. Results from all the SABR screens from the top expanded TCRs ΔFR (y-axis) vs ΔRank plotted for each of the top CD4+ TCR clones screened against the I-Ag7-SABR-II library.

**Fig S6.**
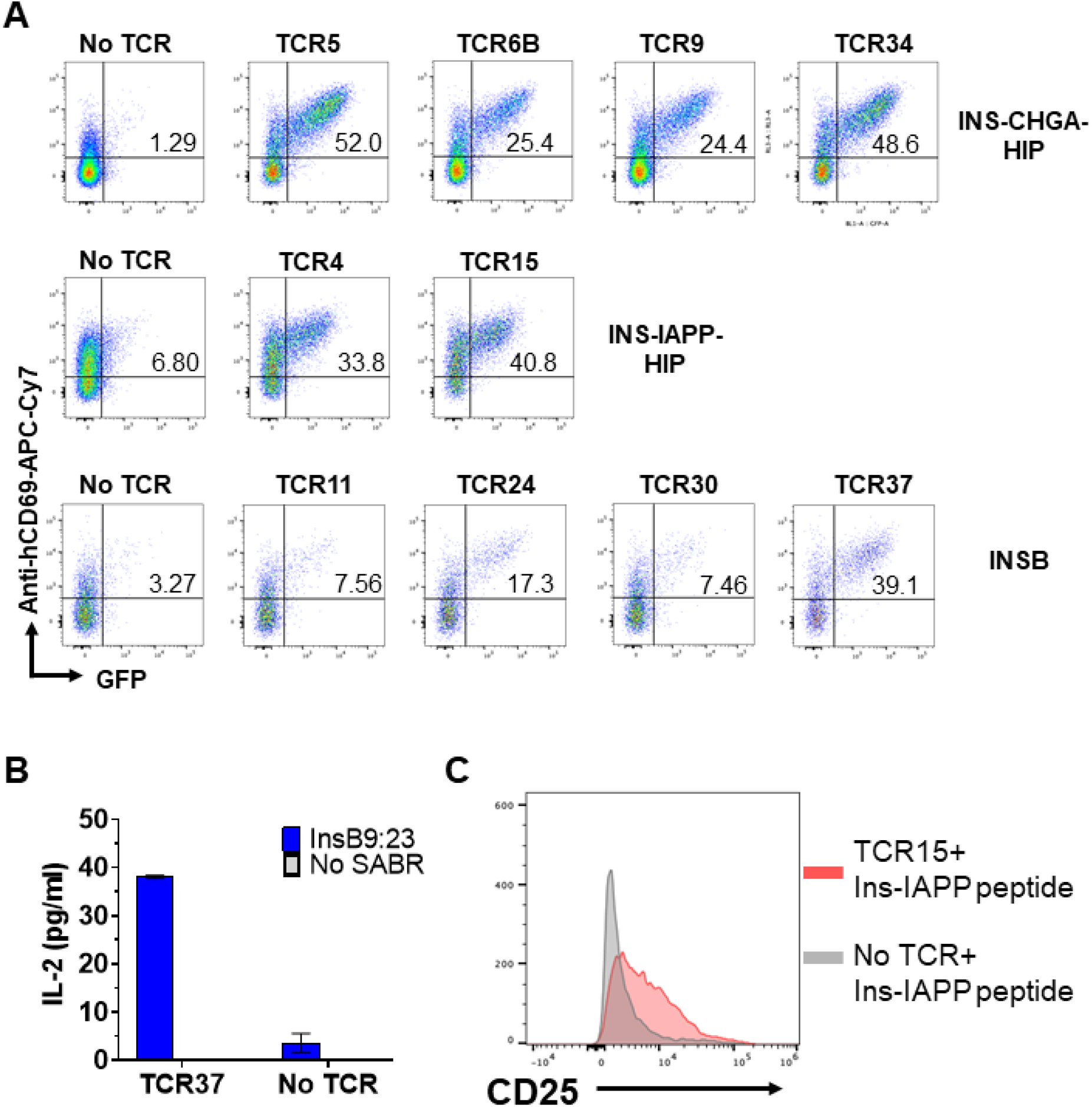
Representative validation of putative hits from the I-Ag7-SABR-II library screens. **A**. Representative flow plots of GFP+CD69+ NFAT-GFP-Jurkat cells expressing single-SABR-II epitopes from co-incubation with TCR expressing Jurkat cells for validation of putative hits found in I-Ag7-SABR-II library screens. The TCRs and their respective cognate epitopes are indicated. **B**. Murine IL-2 ELISA from 24 hr co-incubation of 5KC cells expressing TCR37 with NFAT-GFP-Jurkat cells expressing the InsB9:23 epitope. **C**. CD25 expression measured on primary murine CD4+ T cells expressing TCR15 after 24 hr coincubation with Bone Marrow Dendritic Cells pulsed with either the InsC-IAPP peptide (LQTLALNAARDP) or no peptide.

**Fig S7.**
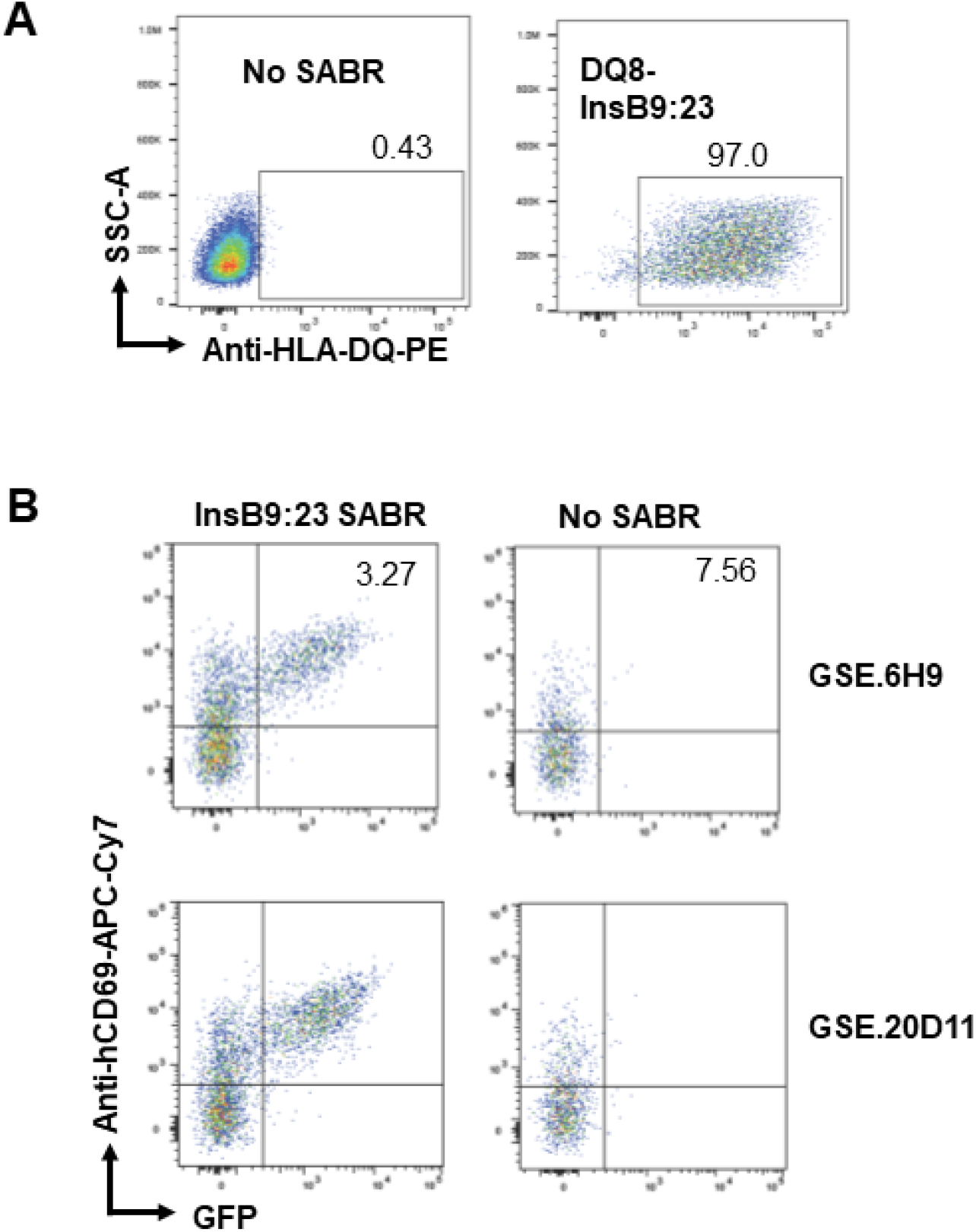
Representative coincubations with HLA-II SABRs. **A**. Representative expression or HLA-DQ8-InsB9:23 SABR-II after transduction of NFAT-GFP-Jurkat cells respectively. **B**. Representative flow cytometry plots of SABR-II expressing GFP+CD69+ NFAT-GFP Jurkat cells after co-incubation with TCR expressing Jurkats. The respective TCRs and SABRs are indicated.

